# The role of Absorbing Markov Chains in childhood cancer

**DOI:** 10.1101/2022.12.12.520113

**Authors:** David H. Margarit, Marcela V. Reale, Ariel F. Scagliotti, Lilia M. Romanelli

## Abstract

Absorbing Markov Chains are an important mathematical tool used for different applications in science. On the other hand, cancer and its metastases in children have a significant impact on health due to their degree of lethality. Therefore, the aim of this work is to model the metastatic pathways of the main childhood cancers worldwide. The probabilities of generating metastases, from a primary site to secondary and tertiary sites, were characterized by constructing a directed graph and the associated transition matrix. In addition, the time of absorption and the probabilities of absorption by each absorbing state were calculated.

## 1. Introduction

Cancer is the deadliest disease in the world according to the World Health Organization (WHO), one in six deaths is due to this disease that afflicts people at any age (children, adults and the elderly)[1]. The determining factors are very varied[2]. In general, cancer is the uncontrolled replication of cells of a tissue, altering its intrinsic properties and damaging it, even causing death[3]. Cancer can not only establish itself in the organ where it originates, but it can spread and colonize other tissues or organs, generating a new one there. This phenomenon is known as metastasis[3]. Although it is estimated from which organ to which other metastasis can be generated, it is not known what these routes will be like when there is subsequent cancer (metastasis of metastases) or when it will end up in a tissue where it will not spread further[3]. In children, childhood cancer seriously affects their health, causing premature deaths and serious side effects in those who manage to recover[4]. It cannot be ignored, and it is part of the motivation of this work, that in paediatric cancer the most common cancers are not the same as in adults[5, 6], as well as the sites of metastasis or the frequency with which they occur.

From a mathematical point of view, Markov Chains promote a tool that is used in different fields due to its potential for predictability based on probabilities that are determined by linear algebra. Markov Chains are used not only in mathematics but also in other science[7, 8, 9]. In this sense, it is possible to extrapolate and use them in the biological systems, in particular, in cancer and metastasis

In previous work, we used Markov Chains for statistics of Argentina[10]. Now, having into account the importance of modelling mathematically routes the metastasis in childhood cancer, we analyse metastatic pathways in the main haematological and solid cancers for children aged 0-14 years. The data used here was obtained from the International Agency for Research on Cancer (IARC)[11], belonging to WHO.

In this way, by using Markov Chains, we will analyse the different steps between the metastases from one organ (primary site) to others, whether they are secondary or tertiary sites. Those organs with a low probability of generating metastases are called absorbing states, therefore, to contemplate this phenomena this work is developed through Absorbing Markov Chains. Thus, we will probabilistically analyse how organs are related to childhood metastases and the role of absorbing states through current information.

## 2. Methodology

There are three general ways to spread cancer cells and generate new cancer in another organ: through the blood circulation, lymph nodes and transcoelomic tissues[12]. Blood vessels are the primary route to spread to distant organs. Meanwhile, lymphatic vessels provide a route to local lymph nodes, and frequently after metastasis, they spread through the blood. Although the spreading through the blood seems to be quite common, the lymphatic spread appears to depend on the location of the primary tumour. For example, cancer of bone and soft tissue (sarcomas) firstly spread through the blood, while kidney cancer spreads through the lymphatic system. Transcoelomic spread is rare and appears to be restricted to mesotheliomas and ovarian carcinomas[13].

As the current WHO-IARC statistics, those cancers that commonly affect children are related to the circulatory and lymphatic systems (leukaemia, Hodgkin’s lymphoma, non-Hodgkin’s lymphoma)[11]. Subsequently, solid tumours are the ones that continuous the list (brain, kidney, liver, ovary, testicular, thyroid, nasopharynx, lip and oral cavity, and bone)[11]. However, bone, liver, and brain are organs that, when they are primary sites, do not generate metastases in other organs (or at least there is a very low probability of this happening)[11, 12], these are called absorbing states or sites. It is worth clarifying that we will not distinguish between cancer subtypes with the intention of being clearer and more global in the analysis. Fig. 1 shows the directed graph and how cancer can migrate from a certain primary cancer to another organ generating a secondary one.

**Figure 1:**
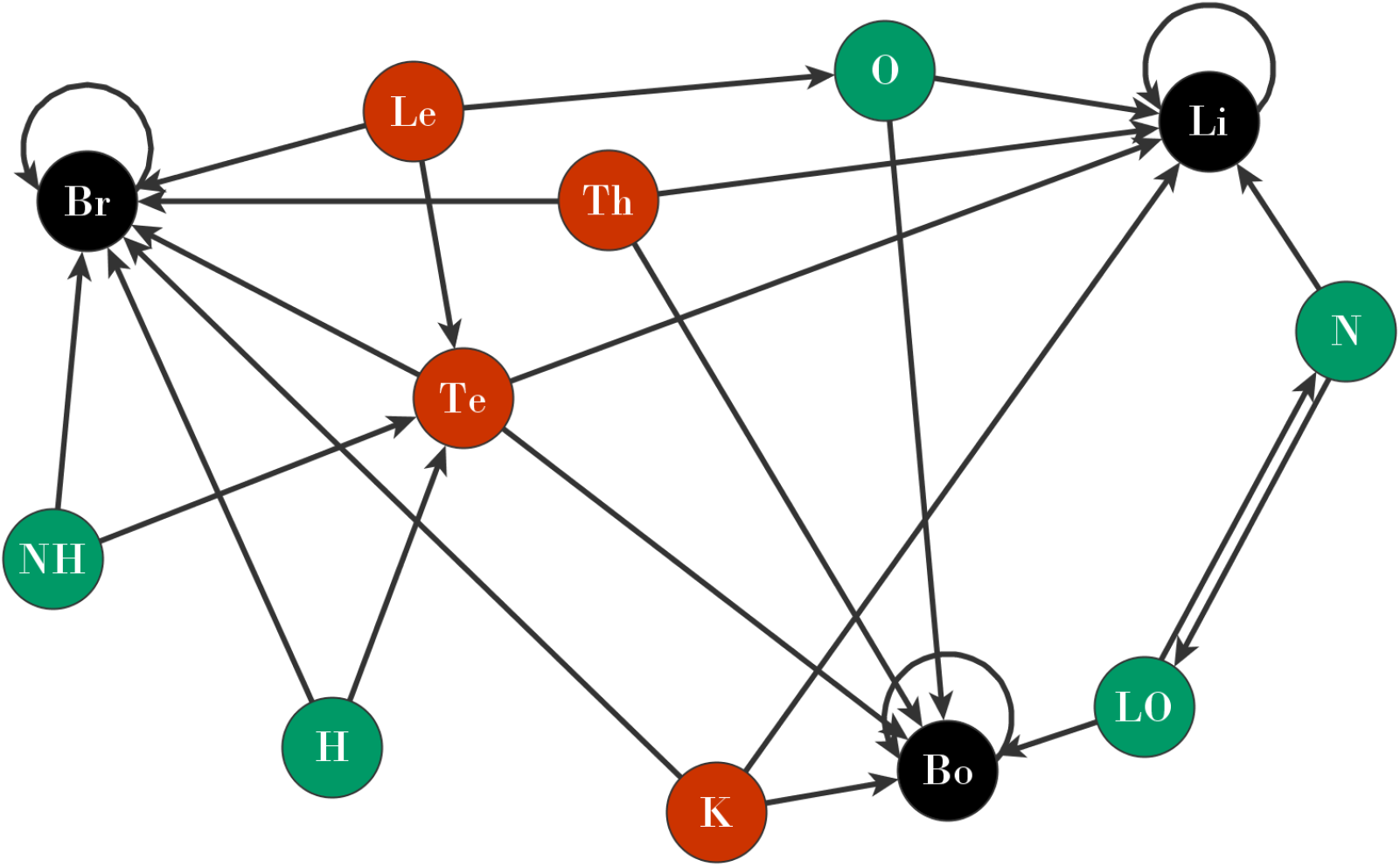
Graph of the routes of metastasis of the most common childhood cancers. Nodes in green: from these organs, metastasis is generated mainly in 2 organs; in red: these organs generate metastasis mainly in 3 organs; finally, in black, the absorbing states, these generally do not metastasise.

Then, from the most reported cancers in children by WHO-IARC statistics[11], we found that these primary sites can generate metastasis in the following sites:

- Leukaemia: ovaries, testicles and brain.[14, 15, 16]
- Hodgkin lymphoma: testicles and brain.[15, 17, 18]
- Non-Hodgkin lymphoma: testicles and brain.[15, 17, 18]
- Kidney: brain, liver and bones.[5, 19, 20]
- Ovaries: liver and bones.[21, 22]
- Testicles: brain, liver and bones.[5, 23, 24]
- Thyroids: brain, liver and bones.[25, 26, 27]
- Nasopharynx: lip/oral cavity and liver.[28, 29]
- Lip/oral cavity: nasopharynx and bones.[30, 31]
- Brain: usually does not metastasize to another organ.
- Liver: usually does not metastasize to another organ.
- Bones: usually does not metastasize to another organ.

In general, brain, liver and bone cancer do not spread to other organs. Therefore, if they generate metastasis is only on themself.[32, 33]

For simplicity, and considering the name of the organs, cancers will be referred to by a symbol as depicted in Table 1.

**Table 1:**
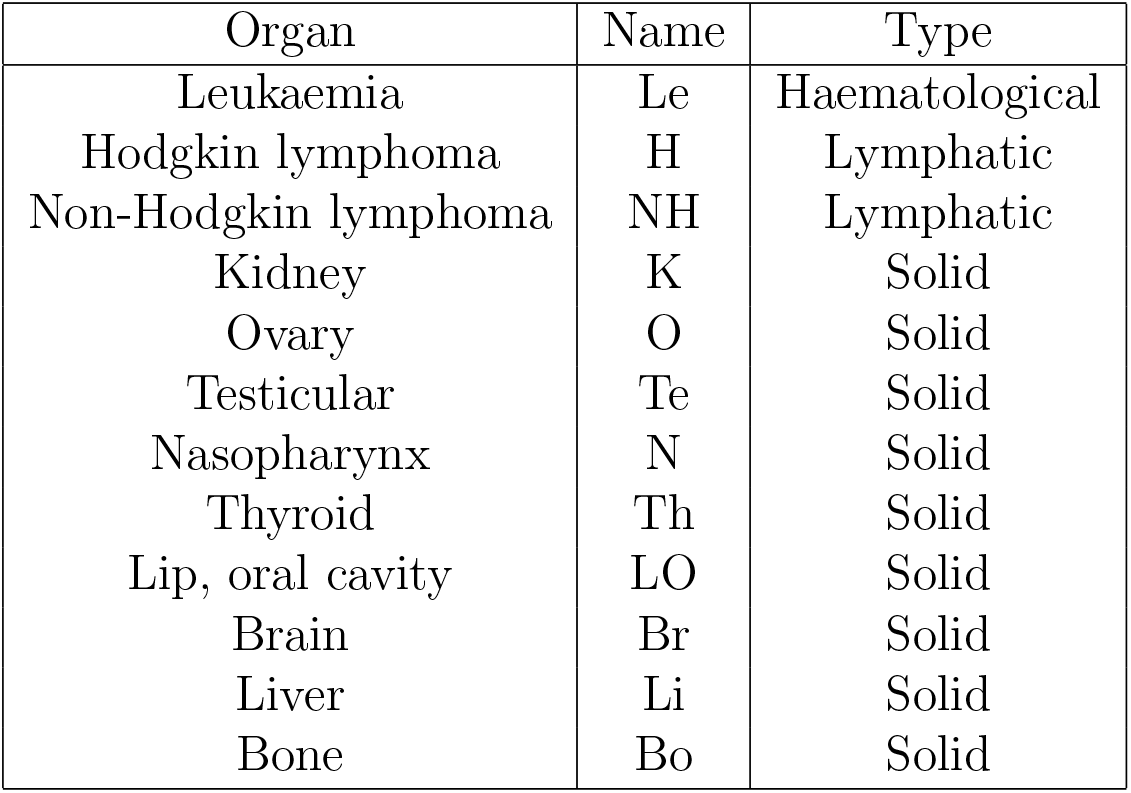
Symbols for organs depending on their type.

In the cancers of Table 1, it does not appear the lung. Even though the lung is known to be the organ where the majority of organs can develop metastasis, this is not the case for paediatric cancer[34, 35, 36].

## 3. Construction of the transition matrix and absorbing states

Let *X*_0_(primary site) be the organ where cancer originated, and *X*_1_ is the state of the process where a new cancer is formed coming from a primary site. We consider it as a step of the Markov matrix when a new tumour has already been found in another organ. Likewise, the same concept applies to the second step matrix (this will be discussed later). In terms of the Markov Chain formalism[37], the above implies that the transition matrix has been built under the assumption of the existence of the tumour in *X*_0_ that will develop metastasis in *X*_1_. Then, the probability of an organ (*i*) develops metastasis into another (*j*) is:

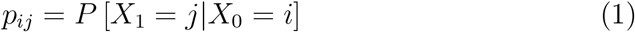

where *i,j* = 1,2,3,…,*m* refer to the *m* = 12 organs taken into account in Table 1.

In other words, *X*_1_ (metastasis from the primary site or a secondary site) will have a probability of being metastasis from some primary site called *X*_0_. If the tertiary site (metastasis from the secondary site) exists, it will be labelled as *X*_2_.

Generalizing these probabilities *p_ij_* for all the *m* cancers, we construct the called transition matrix called *P*.

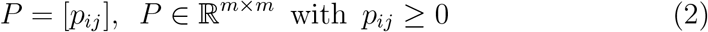

Since there is no general agreement on the probability of metastasis in children, because this differs in the reported cases, we will consider that the sites of metastasis are equiprobable. That is, if an organ *i* has *ρ* possible metastases, the probability *p_ij_* will be 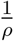.

In terms of the current statistics and data, we built the matrix transition *P* as follows:

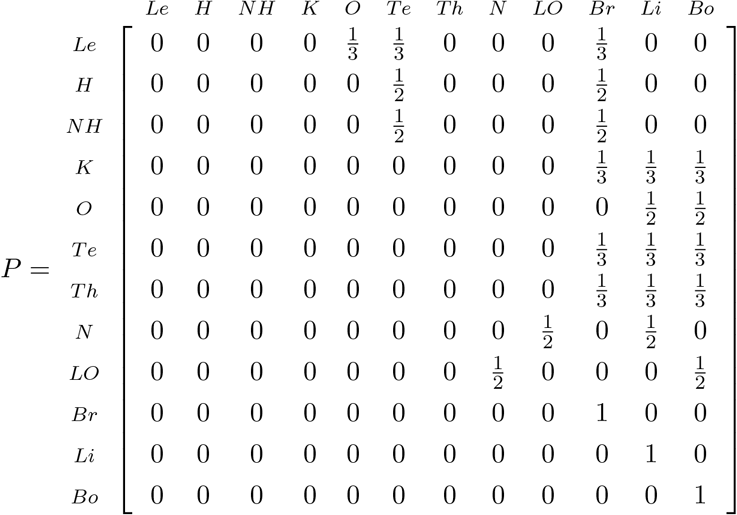

A way to extract information from *P*: in Fig. 2*a* and 2*b* there, are two examples that describe the probabilities of generating metastasis in determinate organs from specific primary sites (thyroids and lip and oral cavity).

**Figure 2:**
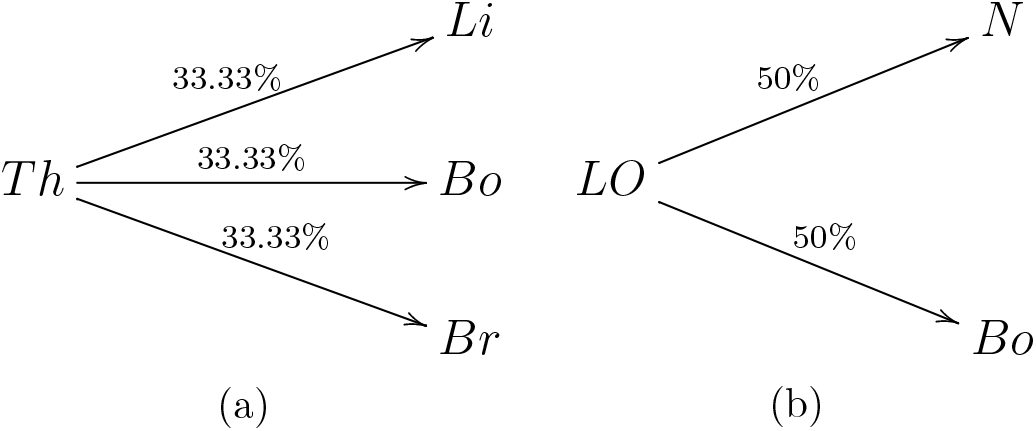
Probabilities of thyroids and lip and oral cavity of generating metastasis given by *P* matrix.

We can observe in the P matrix the existence of Absorbing States in the system. These are states where once they are reached it is impossible to leave[37]. In terms of probability:

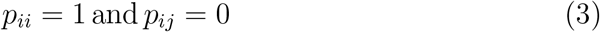

In our system, we can see that in the P matrix there are three absorbing states (the organs brain, liver and bone). In these particular organs, when cancer is originated there, the probability of metastases in another organ is virtually null. However, if the cancer forms elsewhere (first or second), it can metastasise to one of these sites, but no new metastases will form from there. Therefore, it can be considered an absorbing matrix.

## 4. Transition matrix for tertiary sites

A quite important application of Markov Chains is to analyse the probability when a process goes from state *i*, through an intermediate state *k*, to state *j* in two steps[37]. That is:

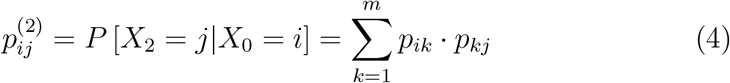

In the context of our models, we can estimate quantitatively the probability of metastasis being generated in a tertiary site, that is, a metastasis of metastases. Generalizing this for all organs, based on Eq. 4, the matrix of probabilities of metastases from metastases (cancer to the tertiary site from the primary site) is given by:

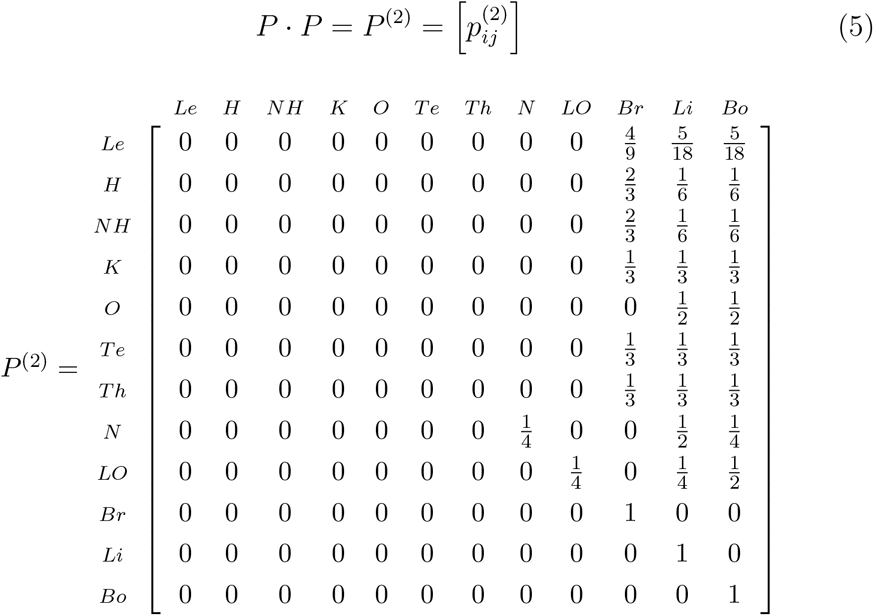

The transition matrix of the second step (*P*^(2)^) shows that, in general, from a primary site there is a higher probability, compared to *P*, of having reached an absorbing state (liver, liver or brain). That is, for metastases of metastases, an absorbing state always has a greater than zero probability of being reached. To quantify this information, such as the number of steps to complete an absorption state, both in general and for a particular organ, we will develop the following section.

## 5. Absorptive capacity by non-absorbing states and number of expected metastases

In Absorbing Markov Chains, the number of steps before the system is absorbed and the probability of absorption by any absorbing state can be found. In order to calculate this information, each transition matrix must be represented in its canonical form[37], called *J*. The canonical *J* matrix is composed by 4 sub-matrices (*N, A, O* and *I_a_*). These smaller matrices contain elements of probability that originate a absorbing states and *n* nonabsorbing states. Thus, there are *a* + *n* = *m* states of the system. For our model, the matrix *J* is represented by:

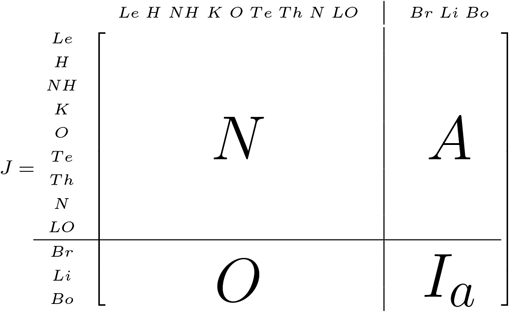

- Sub Matrix *N*: *N* ∈ℝ^*n×n*^ has the probabilities of going from a nonabsorbing state to another non-absorbing state.
- Sub Matrix *A*: *A* ∈ℝ^*n×a*^ has the probabilities of going from a nonabsorbing state to another absorbing state.
- Sub Matrix *O*: *O* ∈ℝ^*a×n*^ represents the probabilities of going from an absorbing state to another non-absorbing state (zeros matrix).
- Sub Matrix *Ia*: *I_a_* ∈ℝ^*a×a*^ this represents the probabilities of remaining inside of an absorbing state (an identity matrix).

Let be fundamental matrix[37] *F* = (*I* – *N*)^-1^, where *I* is an identity matrix with the same dimensions as *N*. Starting in a transient state, the expected number of steps (in our case, a step in the metastasis propagation from one organ to another) before being absorbed by an absorbing state is given by:

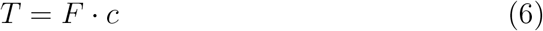

where *c* is a column vector whose entries are ones.

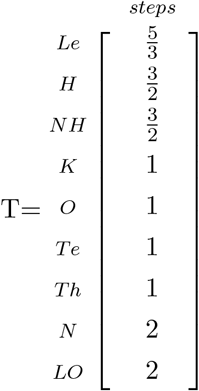

As can be seen, the expected number of steps *T* for metastasis generated in a primary site and ending in an absorbing organ is at most (or close to) 2. Haematological or lymphatic cancers have a number of steps between 1 and 2, which implies that more than one metastasis can occur from the secondary site. Although these cancers do not metastasise in adults (except for very isolated events), in children they can develop a branching metastatic pathway. In the cases of the nasopharynx and lip/oral cavity, they are generally absorbed in the second step of metastasis until they reach one of the three absorbing organs. Finally, the ovaries, the testicles and the thyroid will generate a metastasis and will remain in an absorbing state, something that improves the prevention prognosis.

The vector *T* can be interpreted as a degree of tolerance of the human body. Generally, if the primary sites are not treated, they are known to trigger eventual metastases (in one or more organs), and from there to other sites. Therefore, this information acquired by *T* is an important tool for early diagnosis and potential predictions about cancer sites and pathways depending on their origin.

On the other hand, the probability of absorption of any non-absorbing state by any absorbing states is depicted by the matrix *Z*[37]:

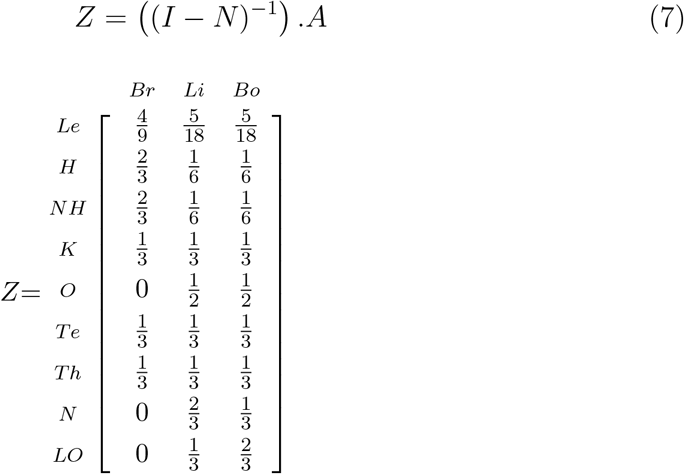

The *Z* matrix has a similar meaning as the vector *T*, except that we observe specifically for each primary site the probability of being absorbed by a certain absorbing organ (states). These results also provide sensitive information for the treatment of childhood cancer. As can be seen from the *Z* matrix, leukaemia and both lymphomas are more likely to end (if they metastasise) in the brain. In the case of the nasopharynx and lip/oral cavity, they finish their route in the liver or in the bones. Finally, the rest of the organs end up being absorbed by any of the three absorbing states. It is important to highlight the difference between the probabilities of absorption and the routes where cancers generated in solid and non-solid tumours can end up.

## 6. Conclusions

Through the use of Absorbing Markov Chains, we were able to quantitatively characterize the metastatic pathways of the main childhood cancers.

It is important to highlight that it was found that in no more than two steps the absorption states (liver, brain, and bones) are reached. In addition, the probabilities of knowing specifically in which absorbing organ the pathway end, depending on the primary site, were characterized.

In the previous section, the matrices T and Z provide relevant information for any preventive treatment in children who are going through this disease.

As a future work, different age groups will be considered in order to compare them, taking into account different age ranges, sex or pathologies related to some types of cancer in particular. Additionally, we will work with a model that implements machine learning to automatically optimize the model when available statistics are updated.

## Acknowledgements

This work is partially financed by PIP - CONICET N^o^11220200100439CO. The authors acknowledge funding from European Union’s Horizon 2020 MSCA-RISE-2016 under grant agreement N^o^734439 (INFERNET Project: New algorithms for inference and optimization from large-scale biological data).

